# The Lambda variant of SARS-CoV-2 has a better chance than the Delta variant to escape vaccines

**DOI:** 10.1101/2021.08.25.457692

**Authors:** Haolin Liu, Pengcheng Wei, Qianqian Zhang, Katja Aviszus, Jared Linderberger, John Yang, Junfeng Liu, Zhongzhou Chen, Hassan Waheed, Lyndon Reynoso, Gregory P. Downey, Stephen K. Frankel, John Kappler, Philippa Marrack, Gongyi Zhang

**Affiliations:** Department of Immunology and Genomic Medicine, National Jewish Health, Denver, CO 80206, USA and Department of Immunology and Microbiology, School of Medicine, Anschutz Medical Center, University of Colorado, Aurora, CO 80216, USA; State Key Laboratory of Agrobiotechnology, College of Biological Sciences, China Agriculture University, Beijing 100193, People’s Republic of China; Department of Biochemistry and Molecular Genetics, School of Medicine, Anschutz Medical Center, University of Colorado, Aurora, CO 80216; Department of Medicine, National Jewish Health, Denver, CO 80206, USA; Department of Pharmacy, National Jewish Health, Denver, CO80206, USA

**Keywords:** SARS-COV-2, COVID-19, Lambda/C.37, Delta/B.1.617, Alpha/B.1.1.7, Beta/ B.1.351, Gamma/P.1, ACE2, RBD, Bamlanivimab, escape

## Abstract

The newly emerging variants of SARS-CoV-2 from India (Delta variant) and South America (Lambda variant) have led to a higher infection rate of either vaccinated or unvaccinated people. We found that sera from Pfizer-BioNTech vaccine remain high reactivity toward the receptor binding domain (RBD) of Delta variant while it drops dramatically toward that of Lambda variant. Interestingly, the overall titer of antibodies of Pfizer-BioNTech vaccinated individuals drops 3-fold after 6 months, which could be one of major reasons for breakthrough infections, emphasizing the importance of potential third boost shot. While a therapeutic antibody, Bamlanivimab, decreases binding affinity to Delta variant by ~20 fold, it fully lost binding to Lambda variant. Structural modeling of complexes of RBD with human receptor, Angiotensin Converting Enzyme 2 (ACE2), and Bamlanivimab suggest the potential basis of the change of binding. The data suggest possible danger and a potential surge of Lambda variant in near future.

---

The SARS-CoV-2 virus has infected over two hundred million people (COVID-19 patients) and caused more than four million deaths to date ^1^. The number of affected people continues to grow rapidly, emphasizing the importance of the rapid use of effective vaccines. Although two Spike mRNA (Pfizer-BioNTech COVID-19 Vaccine and MODERNA respectively) based vaccines and others have been approved for emergency use in the USA ^2,3^, the increasing number of Spike variants that have appeared around the world raise concerns about the continued efficacy of the vaccines ^4^. It has been reported that above 90% of broadly neutralizing anti-SARS-CoV-2 antibodies from COVID-19 patients as well as vaccinated individuals engage in the receptor binding domain (RBD) of the virus Spike protein ^5,6^. Monoclonal antibodies specifically targeting the native form of the Spike developed by different companies have been approved by the FDA for emergency use ^7–10^. An N501Y variant of SARS-CoV-2 (Alpha/B.1.1.7), first emerging in the United Kingdom and spreading to the rest of the world last year, appears much more contagious than the original version ^4^. We found that a single mutation of N501Y confers an ~10 times fold increase of affinity between RBD and ACE2 ^11^. However, this mutation does not affect its binding to the therapeutic antibody, Bamlanivimab ^11^. We concluded that the increase of high binding affinity may account for the high infection rate of the United Kingdom variant while both vaccines and the therapeutic antibody Bamlanivimab should remain their efficacy to combat this newly emerging variant ^11^. However, the same N501Y mutation is also found in a variant (Beta/B.1.351) with mutations of K417N, E484K, and N501Y from South Africa and a variant (Gamma/P1) with K417T, E484K, and N501Y from Brazil ^4^. We found that one additional mutation, E484K, a critical residue involved in the interactions between RBD and Bamlanivimab, completely abolishes the binding between RBD and Bamlanivimab though with no effect of its binding to ACE2 ^12^. It has been reported that a COVID-19 patient was infected a second time by the new variant with E484K mutation in Brazil ^13^ and the Gamma variant (P.1) besides the Lambda variant (C.37) becomes one of the major variants in COVID-19 patients from Argentina and Chile recently in populations with or without vaccines ^14^. It is likely that this critical mutation at E484 to K484 from both the South Africa and Brazilian variants is responsible for the breakthrough infection of the virus in South America. Conversely, although without the dramatically binding affinity enhancing mutation of N501Y in the Delta variant, which first spread in India, it becomes dominated all over the world recently. The Delta variant has two or three mutations within RBD, L452R, and T478K, and some sub-variants with E484Q, which are close but not involve in the direct interactions with ACE2, suggesting minor effects on binding affinity, but it breaks through the protection from vaccines and can infect vaccinated people all over the world ^15–17^. Similar to the Delta variant, the Lambda variant (C.37) does not contain the N501Y mutation either but also with two mutations within the RBD, L452Q, and F490S. However, it becomes dominant in South America and infected vaccinated people ^14^.

These two variants raised major concerns of efficacy of current vaccines and potential coming risks of surge of the virus. Here we present data suggesting that the Lambda variant could be more dangerous than Delta variant and it has a high potential to escape the current vaccines.

## Results

As we reported earlier, N501Y-RBD (N501 mutated to Y501) derived from the United Kingdom variant has a ~10 fold increased binding affinity (0.566nM) toward ACE2 compared to the wildtype (5.768 nM) (**Table 1**) ^11^. It was reported that both the South African variant and the Brazilian variant containing the same N501Y mutation are also highly contagious as that of the United Kingdom variant ^18^. However, they gained two additional mutations, K417N and E484K existing in the South Africa variant, and K417T and E484K in the Brazilian variant. We found that the shared mutation within these wo variants, E484K, completely abolished the binding of the antibody, Bamlanivimab ^12^, suggesting that the area around E484 within the RBD is a hot area to produce broadly neutralizing antibodies. From these previous results, we conclude that SARS-CoV-2 could gain infectivity and escaping through two unique mechanisms: mutations within the receptor binding region to increase the binding affinity toward the receptor, ACE2; mutations within broadly neutralizing antibodies producing regions to escape the immune system of the host. It is likely that both Delta and Lambda variants may utilize the similar strategies to increase their infectivity and break through the immune protection generated by vaccines.

**Table 1.** The binding affinities of variant RBDs to ACE2 and Bamlanivimab.

| Strains | Geographic emergence | Spike RBD mutations | ACE2 binding affinity | Bamlanivimab binding affinity |
| --- | --- | --- | --- | --- |
| B.1 | Wuhan & Europe | Wildtype | 5.765 nM, **8.3 nM** | 0.874nM, **1.4 nM** |
| B.1.1.7/Alpha | England | N501Y | 0.566nM | 0.801nM |
| B.1.351/Beta | South Africa | K417N, E484K, N501Y | 2.935nM | None |
| P.1/Gamma | Brazil | K417T, E484K, N501Y | ~3.00nM | None |
| B.1.617/Delta | India | L452R, T478K | **4.0nM** | **31 nM** |
| C.37/Lambda | South America | L452Q, F490S | **7.6nM** | **None** |
| SARS | China | numerous | **46.6nM** | unavailable |

The Delta variant brings a new surge in USA and all over the world ^15^. Most surprisingly, this variant breaks through the immunity generated by vaccines ^15,16^. However, to people’s relief, it was found that the virus was cleared quickly in the vaccinated individuals ^17^. It is of great interest to understand the potential underlying mechanism of the escaping of this variant. R452/K478-RBD from Delta variant was introduced and expressed in 293F cells as previously reported (Fig. S1) ^11^. Purified protein R452/K478-RBD was subjected to binding assays to ACE2 on Biacore machine. Interestingly, the binding affinity (4.0 nM) (Fig. 1A) between R452/K478-RBD and ACE2 is ~2 times higher than that of wildtype RBD and ACE2 (8.3 nM) (Fig. 1B). From this result, the two mutations within the Delta variant partially explains the high infectivity. At the same time, we tested R452/K478-RBD binding to the neutralizing antibody, Bamlanivimab.

**Figure 1.**
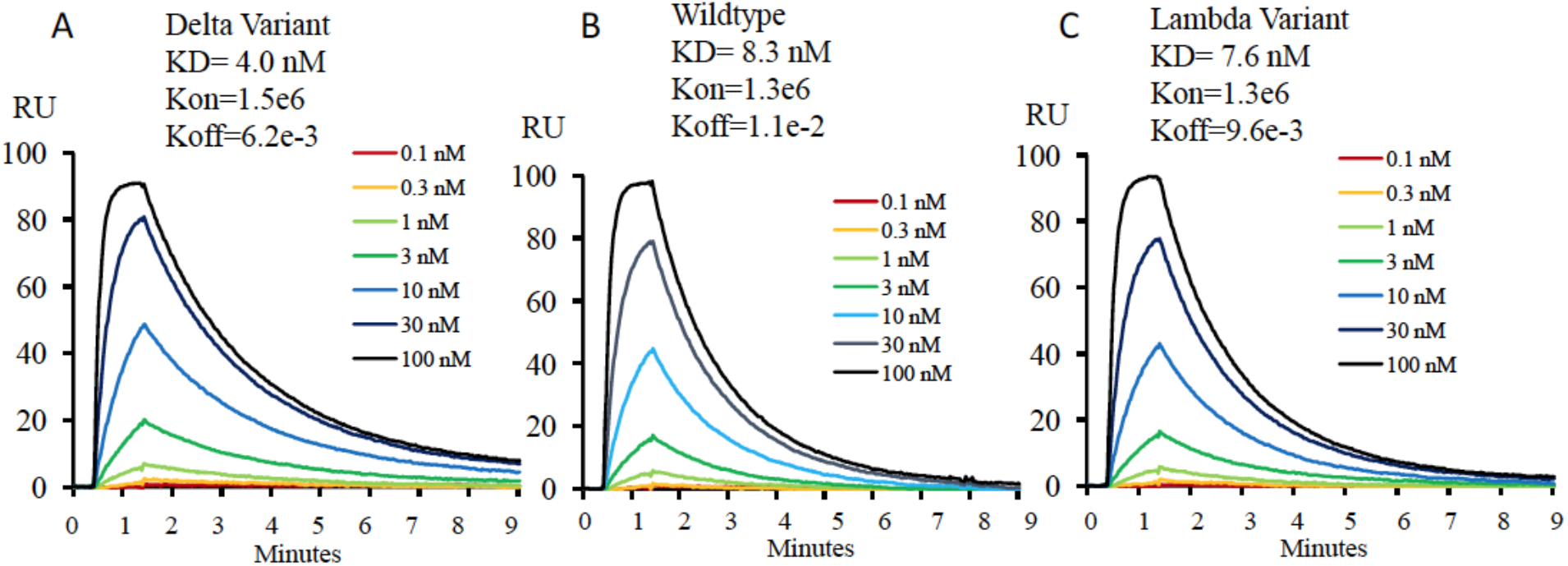
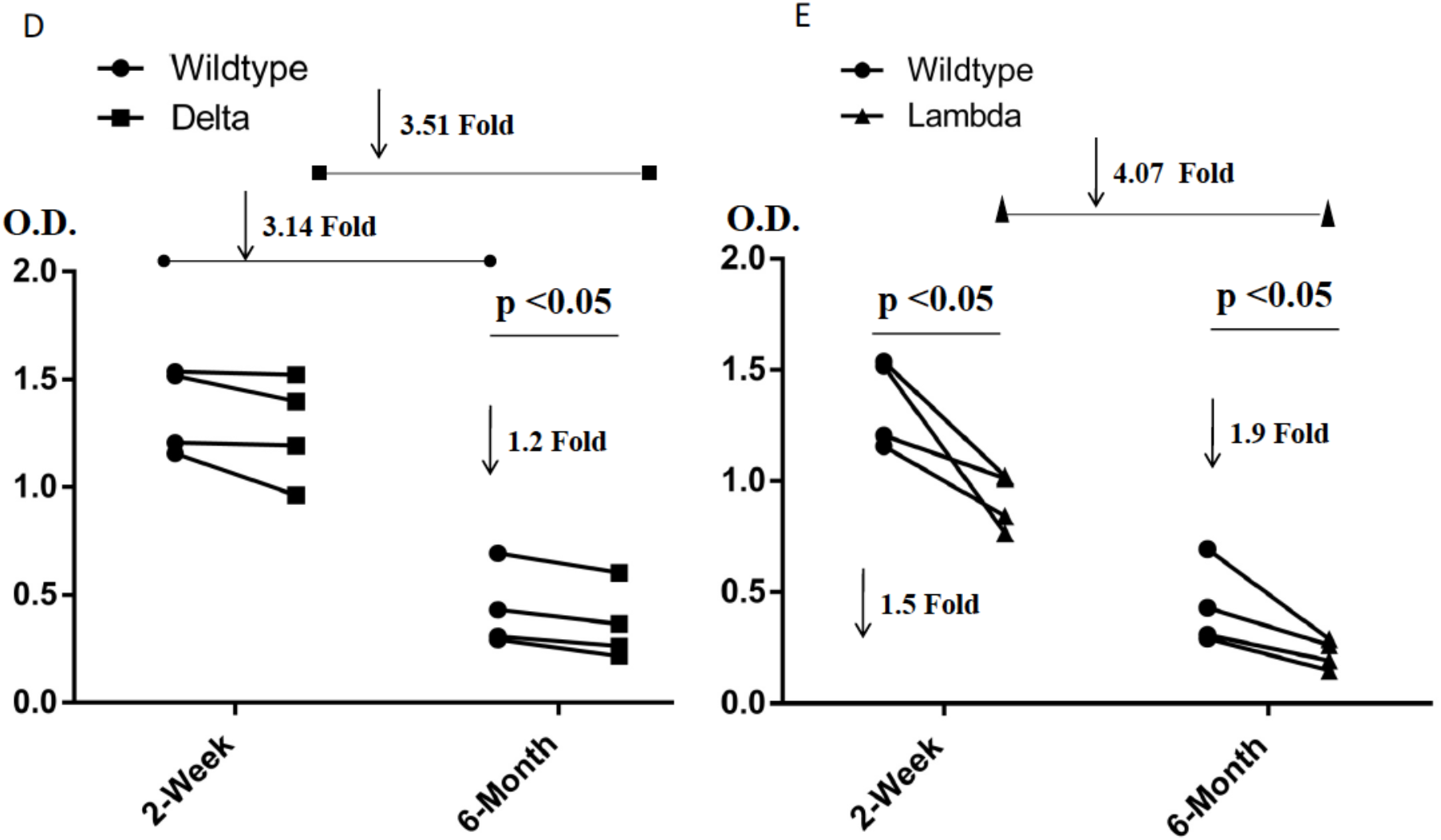
The affinity measurement of the mutated RBDs with ACE2, Bamlanivimab, and Sera from four Pfizer-BioNTech vaccinated individuals. **1A,1B, and 1C**.The binding properties of variant-RBDs to ACE2. A. The binding affinity of the Delta-RBD to ACE2. B. The binding of affinity of the wildtype-RBD to ACE2. C. The binding affinity of the Lambda-RBD to ACE2. **1D and 1E**. The binding properties of variant-RBDs to Sera. D. The binding affinity of the Delta-RBD to Sera. E. The binding of affinity of the Lambda-RBD to Sera.

As expected, compared to wildtype RBD, the binding of R452/K478-RBD to the antibody drops dramatically from 1.4nM to 31.0 nM (**Table 1**). It is expected that an additional mutaion of E484Q within some Delta sub variants could further compomise the binding bewteen RBD and Bamlanivimab. However, the Delta variant RBD still efficiently binds antibodies from sera of vaccinated individuals by Pfizer-BioNTech collected at two weeks after boost, and there is about 1.2 fold drop for sera collected from the same individuals after six months (Fig. 1D). At six months after boost, the antibody against wildtype RBD drops 3.1 fold on average. These data indicate that the Pfizer-BioNTech vaccine induced antibody response is highly effective against the Delta variant. As there is less antibody in the sera collected after six months, the difference of antibody binding between Delta variant and wildtype RBD would become more obvious. It has been reported that the lower serum antibody level after vaccination for a period of time was correlated with breakthrough infection ^19^, and hence a third boost could be necessary to keep serum antibody at a higher level. To further understand the molecular basis of the interactions, the two mutations within the RBD are investigated through structural modeling based on existing structures of the wildtype RBD with ACE2 or antibody. Consistent with the binding data, the T478K mutation may bring an additional hydrogen bond between RBD and ACE2 (Fig. 2A), it may partially explain the ~2 fold increase of the binding (Fig. 1A & B). In Bamlanivimab binding, the L452R mutation is located within an originally hydrophobic interaction interface between RBD and Bamlanivimab, and therefore, the introduction of the positively charged sidechain of arginine of the mutation completely disrupts the hydrophobic interface (Fig. 2B). This explains the loss of function of antibodies including Bamlanivimab targeting this hot area and causing the escape of the variant.

**Figure 2.**
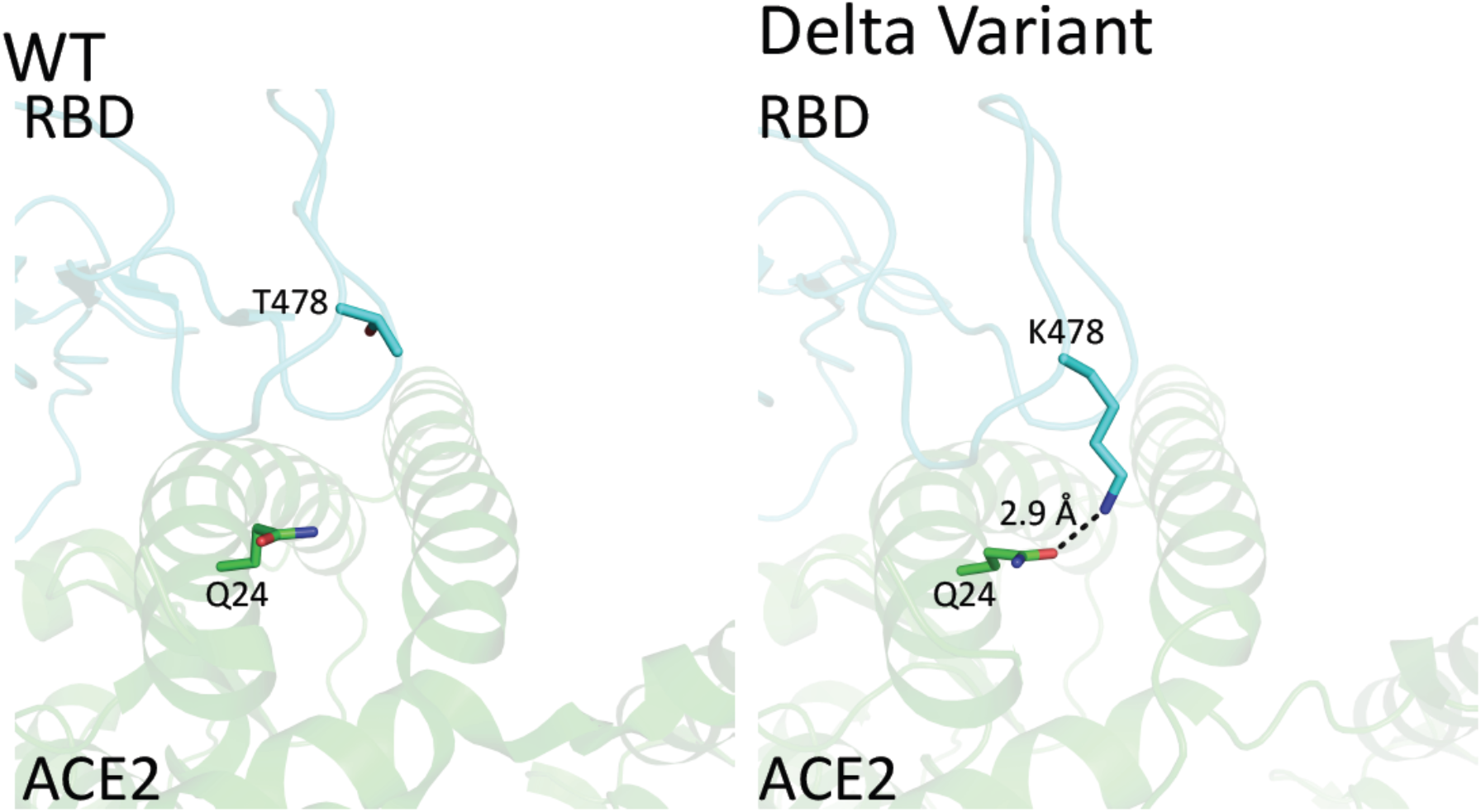
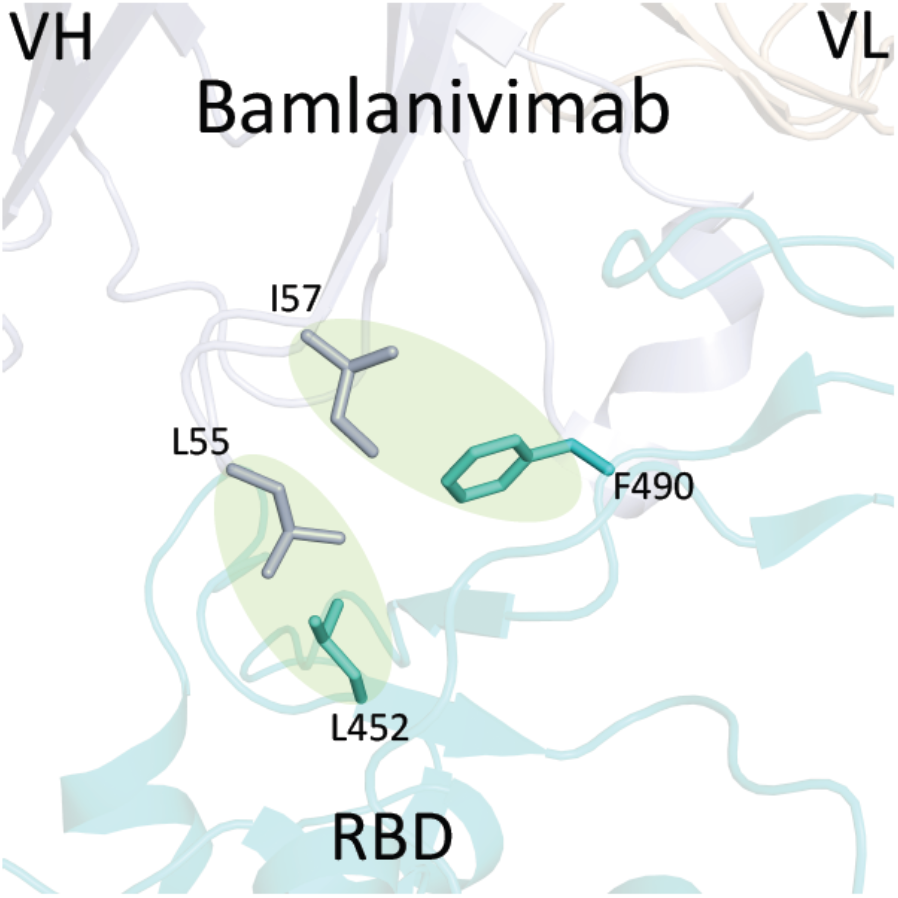
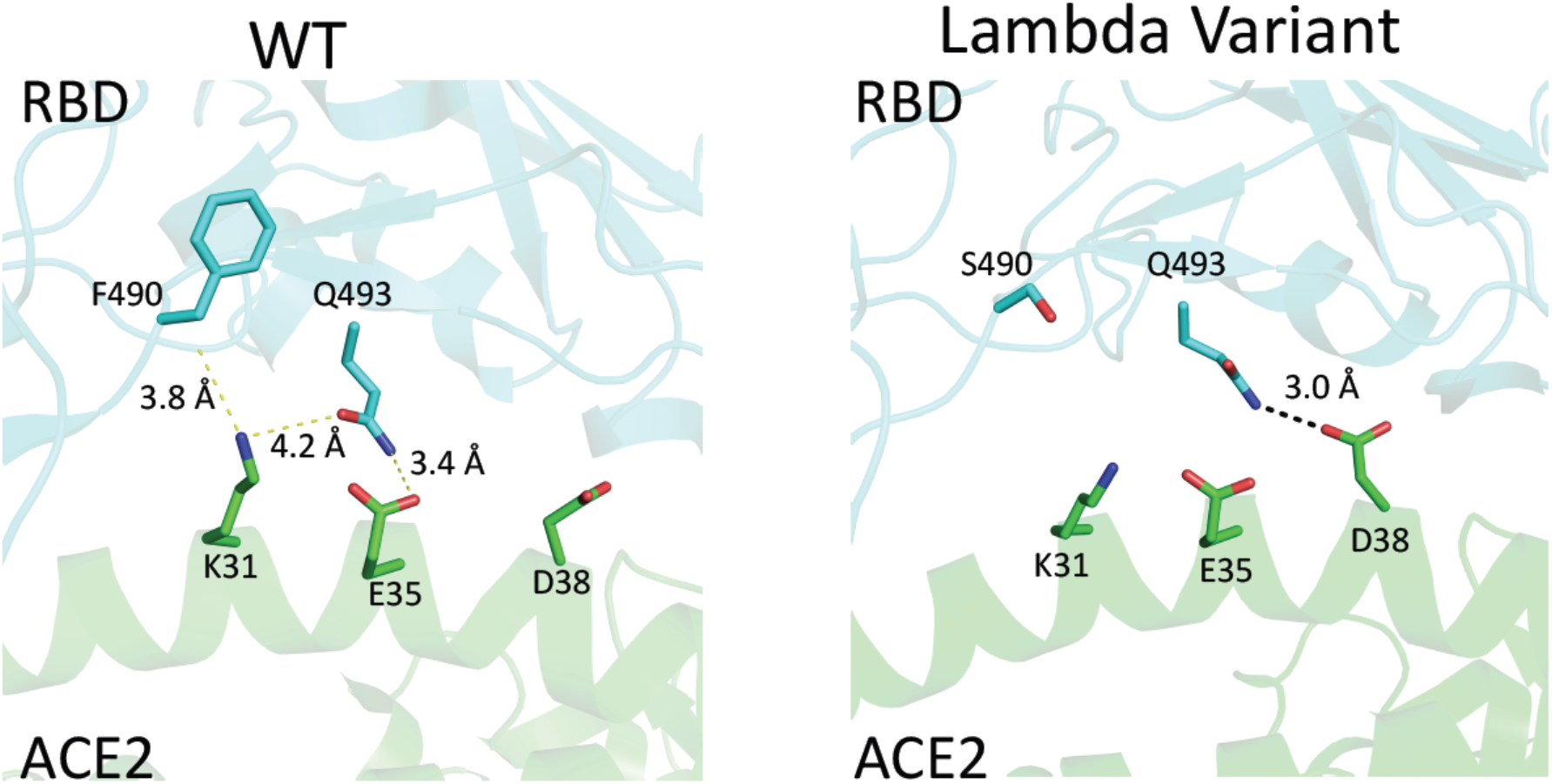
The structural modeling between the mutated RBDs with ACE2 and the therapeutic antibody, Bamlanivimab. **2A.** The mutation of RBD within the Delta variant that is close to the ACE2 binding site. A. T478 of wildtype RBD does not make any interaction with ACE2. B. T478K mutation of Delta-RBD could introduce an additional hydrogen bond between the sidechain of K478 from RBD and the sidechain of Q24 from ACE2. **2B.** The mutations of RBD within both Delta variant and Lambda variant that are close to the Bamlanivimab binding site. The L452R mutation within Delta could disrupt the hydrophobic core formed between RBD and Bamlanivimab due to introduction of the positive charge of R452. Similarly, mutations of L452Q and F49OS within Lambda variant brings polar sidechains to disrupt the hydrophobic core of interaction interface between RBD and Bamlanivimab. **2C.** The mutation of RBD within the Lambda variant that is close to the ACE2 binding site. A. F49O of wildtype RBD barely makes any interactions with ACE2. B. S49O mutation of Lambda-RBD does not introduce any obvious interactions either though Q493 from RBD and D38 from ACE2 may form a weak hydrogen bond.

The Lambda variant emerged in South America later last year and becomes dominant in some South American countries and starts to spread to the world recently ^14^. It seems easily escape the defensive immunity by vaccines ^14^, which raises major concerns of possibility of this variant could be a next surging candidate spreading in the world. Similar to that of the Delta variant, we obtained the protein of 452Q/490S-RBD from 293F cells (Fig. S1). The purified protein is subjected to binding assays toward ACE2, Bamlanivimab, and Sera from vaccinated individuals. Interestingly, 452Q/490S-RBD has a similar binding affinity toward host receptor ACE2 (7.6 nM to 8.3 nM) (Fig. 1B & 1C). However, it is found that the Lambda variant RBD completely loses the binding toward Bamlanivimab (**Table 1**). Without surprise, the ability of this variant RBD to bind antibodies from sera of vaccinated individuals drops dramatically, ~1.5 fold from sera collected at two weeks after boost while ~1.9 fold from sera collected at six months (Fig. 1E). This result raises the alarm of the high potential of breakthrough and escaping of this variant from current vaccines either originated from inactivated viruses, which failed to protect vaccinated people in Chile and Peru as reported ^14^ and possibly do the same to people vaccinated by vaccine from Pfizer-BioNTech or others all over the world. To further understand the molecular basis, the mutations are mapped to the structure of RBD with ACE2 or Bamlanivimab. Consistent with binding data, the mutation of F490S is located close but outside the binding region between RBD and ACE2 (Fig. 2C). In Bamlanivimab binding, this mutation is also located to the hydrophobic interaction interface between RBD and Bamlanivimab (Fig. 2B), joined by the mutation of L452Q (Fig. 2B), which brings polar features into the hydrophobic core and weakens the hydrophobic interactions, accounting for the complete loss of the binding of RBD to Bamlanivimab.

One interesting observation from the experiments is that the overall antibodies from Pfizer-BioNTech vaccinated individuals drop dramatically (~3.5 times fold drop for the Delta variant, Fig. 1D, and ~4.0 times fold for the Lambda variant, Fig. 1E) after six months. This could be one of the reasons that even vaccinated people could get infected. In this regard, these data suggest that the Lambda variant has a better chance than the Delta variant to escape the protection built up by different vaccines.

## Discussion

The original wildtype of SAR-CoV-2, which initiates the outbreak of the pandemic at Wuhan, China, is featured with ~6 times fold higher binding affinity of RBD and ACE2 than that of SARS-CoV-1 (8.3 nM to 46.6 nM, **Table 1**) along with an obtaining of a furin cutting site. Similar to the wildtype, every widely spreading strain owns some unique features to dominate at some places or times. The Alpha strain with an N501Y mutation within RBD, also existing within both Beta and Gamma variants and an additional E484K mutation, were dominant pathogens in United Kingdom, South Africa, and Brazil respectively. Interestingly, both Alpha strain and Gamma strain are still major contributors in Argentina currently while the Delta variant and the Lambda variant took over and become major threats to human beings currently ^14,16^. Most importantly, besides the Gamma variant, both Delta and Lambda variants also could escape the immunity generated from vaccines, which become challenges for people to completely contain the virus and go back to normal life all over the world. Understanding the underlying mechanisms which trigger the higher infectivity and breakthrough of the variants are essential for us to battle the virus. We addressed the potential basis of high infectivity of Alpha, Beta, and Gamma variants earlier and found that the mutation of N501Y within the RBD domain is critical for the high infectivity of all three variant strains while the E484K mutation within both the Beta and Gamma variants, which abolished the binding to the therapeutic antibody, Bamlanivimab, could be the reason to escape the protection built up from previous infections or vaccines ^11,12^. In this report, we dissected the potential factors that confer the properties of high infectivity and immune resistance within both the Delta and Lambda variants and further suggest that the Lambda variant could be a major player in future surges.

From our biochemical analysis, the Delta variant does increase the binding to ACE2 slightly compared to the wildtype. The mutation of L452R could lead to the decrease of its binding to the therapeutic antibody, Bamlanivimab. However, the mutated RBD still binds to most of the antibodies generated from Pfizer-BioNTech vaccine. This is consistent with the report that Pfizer-BioNTech vaccinated people contracting the Delta virus cleared the virus quickly ^17^, suggesting the protection from the vaccine even with the escaping of the virus ^15^. This is also obvious that above 90% COVID-19 patients infected with the Delta variant in USA were not vaccinated currently ^20^, a clear and assured signal that vaccines protect people. However, our data could not explain the fast accumulation or much higher transmissibility of the Delta variant in patients. A recent research finds out that the mutation of P681R outside of RBD, which enhances the furin cut site, is responsible for the high infection ability ^21^. This reminds us the obtaining of the furin site within the wildtype SARS-CoV-2 compared to SARS-CoV-1 ^22^ along with much higher binding affinity of RBD and ACE2 for SARS-CoV-2 triggered the pandemic ^23^, and it also suggests that the furin cutting site could be gained through natural selection. Based on the accumulating data, we attempt to derive that the gain of the furin site and the strength of the binding between RBD and ACE2 could account for the infection ability of the virus.

The story of the Lambda variant could tell us the underlying mechanism of the escaping of the virus. First, the mutations of L452Q and F490S do not bring any enhance of the binding between RBD and ACE2. However, both mutations of F490S and L451Q lead to the disruptions of a hydrophobic patch, which is a critical area for the binding of broadly neutralizing antibodies. This is confirmed by antibody binding assays. The mutated RBD from the Lambda variant decreases the ability to recognize broadly neutralizing antibodies, and this explains why vaccinated people in Peru, who are vaccinated with inactivated viruses, still got infected by this variant. Our data suggest that sera from individuals vaccinated with Pfizer-BioNTech vaccines could be compromised by this variant as well. A recent report derived from the study of a pseudo-virus suggests that mutations of T76I and L452Q lead to the higher infectivity of the Lambda variant while a 7-amino-acid depletion within the N-terminal domain (outside RBD domain) accounts for the immune resistance (or escaping) ^24^. However, T76I is out of the RBD domain while L452Q is out of the ACE2 binding motif of RBD. It is true that the N-terminal of the Spike does generate some broadly neutralizing antibodies, but it is very limited ^5,6^.

Furthermore, there are some shorter depletions of the N-terminal domain within other variants as well. Interestingly, the titers of broadly neutralizing antibodies dropped quickly for vaccinated people after six months, suggesting that the protection of vaccines could weaken with time. From the above analysis, the major factor that confers the breakthrough of the Lambda could be due to obtaining novel antigenic property to circumvent the recognition of broadly neutralizing antibodies generated from vaccines, which could be enhanced by the weakening of protection of the vaccines with time. In this regard, the lambda variant has a better chance to escape the protection from current vaccines than that of the Delta variant and could be the next major pathogen.

It has been almost two years since the breakout of the virus since the end of 2019, but it is still not foreseeable when human beings will completely control the virus, though therapeutic drugs and vaccines have played critical roles to suppress and slow down the spreading of the virus. It becomes clear that the furin site and high binding affinity between RBD and ACE2 are responsible for the high infectivity while mutations mostly within the RBD accounting for the escaping of the virus. One intriguing question is why above 90% broadly neutralizing antibodies either from COVID-19 patients or vaccinated people are against only RBD of the spike ^5,6^. The production of broadly neutralizing antibodies from B Cells is regulated by T cells, and it suggests that the protecting T cells from COVID-19 patients or vaccinated people are likely limited to the RBD portion as well. One most possible explanation is that the four disulfide bridges within RBD, which stabilize the three-dimension structure of RBD, confer the domain as a dominant antigen responsible for host immune responses. Learning from this fact, it could be a promising strategy to introduce artificial disulfide bridges into some conserved regions within the spike protein, such as the proximate membrane fusion region, the N-terminal region, etc, to expand usage of antigens of the entire spike protein for immune response to trigger wider production of broadly neutralizing antibodies as well as T cell protection.

## Methods

### SARS-CoV-2 RBD plasmid cloning and protein expression

SARS-CoV-2 wildtype RBD (319-541aa) was cloned to pCDNA3.1 vector with 6 histidine tag. The Delta variant RBD with L452R and T478K mutations was created by quick change mutagenesis. The Lambda variant RBD with L452Q and F490S mutations was made in the same way. All plasmid constructs were verified by DNA sequencing. RBD proteins were expressed by transient transfection of 293F cell. RBD proteins were purified using nickel column and further purified by Superdex-200 Gel-filtration size column.

### ACE2 protein biotinylation

Human ACE2 (1-615) with BirA tag was cloned to pCDNA3.1 and expressed in 293F cell. Biotinylation of BirA tag was carried out by BirA enzyme. The biotinylated ACE2 was further purified by Superdex-200 Gel-filtration size column.

### Affinity measurement of RBD binding to ACE2

The affinity was measured in a Biacore 3000 machine. Biotinylated ACE2 was coated on the streptavidin chip. A series dilutions of wildtype or mutant RBD were injected at 20 μl/min for 1 minute and dissociated for 9 minutes. The affinity was calculated by BIA evaluation software.

### Affinity measurement of RBD binding to Bamlanivimab

The affinity was measured in a Biacore 3000 machine. Bamlanivimab was coated on CM5 chip. A series dilutions of wildtype or mutant RBD were injected at 20 μl/min for 1 minute and dissociated for 5 minutes. The unbound RBD was eluted by 10mM glycine pH 1.7. The affinity was calculated by BIA evaluation software.

### ELISA measurement of serum RBD antibody from vaccinated donors

Wildtype, Delta or Lambda variant RBDs were coated on the same ELISA plate at 20 μg/ml overnight in cold room. The plates were washed and blocked with PBS with 30% FBS.

Human sera at 2-week after mRNA vaccine boost and 6-month after boost from four healthy donors were used. The sera were diluted at 1:100 or 1:1000. Same volume of diluted serum sample was incubated with wildtype, Delta or Lambda variant RBD coated wells. The bound antibody was detected by AP conjugated goat- anti- human IgG Fc specific antibody.

### Protein docking and binding affinity prediction for mutants

Protein docking studies were performed by HADDock 2.2 server ^25^. The crystal structure of SARS-CoV-2 RBD bound with ACE2 (PDB code: 6LZG) was selected for docking. Delta variant (L452R_T478K) and Lambda variant (L452Q_F490S) structures were prepared by Coot software ^26^. One of the conformations from alternative residues in the structure was deleted manually to meet the docking server criteria. The major contact residues in 6LZG (hydrogen bond and salt bridge) were selected as restrained residues for docking (Q24, E35, E37, D38, Y41, Q42, and Y83 for ACE2; N487, Q493, Y449, and Y505 for RBD). The best model from the docking was selected for the next analysis. All protein structural figures are prepared by PyMOL.

## Acknowledgements

We thank National Jewish Health for supports and people in the Kappler/Marrack groups for help. H.L. is partially supported by NIH grant (GM135421 to G.Z,) and NB Life Laboratory LLC. Biophysics Core at Anschutz Medical Campus for Biacore binding assays. We also thank the Colorado Department of Public Health and Environment (CDPHE) for authorization to use the residual therapeutic antibodies after COVID-19 patient infusions.

## Contributions

H.L., P.M., and G.Z. for designing; H.L. for most experiments; P.W., Q.Z., J. L., J.Y., J.W., N.J., J.L., Z.C., K.A., W.D., S.P., H.W., C.J., L.R., G.D., S.F., and J.K. for some experiments; H.L., P.M., G.Z. for final data analysis and writing up.

## Competing interests

H.L. is partially supported by NB Life Laboratory LLC, G.Z. holds equity at NB Life Laboratory LLC. We do not have any financial relations with Pfizer-BioNTech, Moderna, Eli Lilly, or Regeneron.

## Note

The binding affinity constants in Black were generated from our previous experiments ^11,12^. The binding affinity constants in Red are generated from current experiments. Due to the batch difference and different Biacore machines involved, we have retested the binding affinity between wildtype RBDs to ACE2 and Bamlanivimab. For example, the binding affinity between wildtype RBD and ACE2 was 5.75 nM in an old Biacore machine, but 8.3 nM in a new Biacore machine. Similarly, the binding affinity between wildtype RBD and Bamlanivimab was 0.874 nM in an old Biacore machine while it is 1.4 nM in a new Biacore machine.

## Supplementary Figure

**Figure S1.**
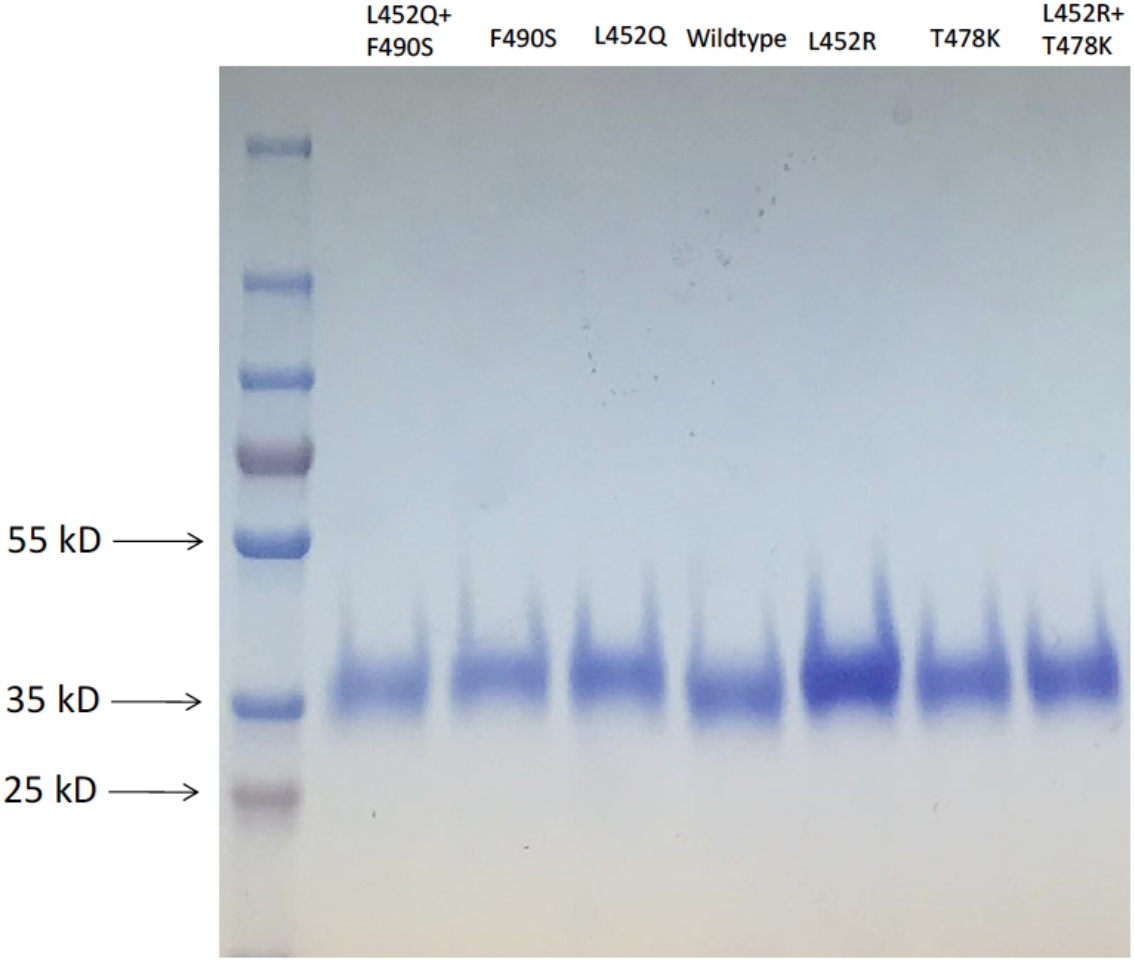
RBDs with different mutations were expressed in 293F cells and purified to high purity for further binding assays.

